# Understanding PebS–Ferredoxin Recognition: A Structural Perspective on Viral and Host Redox Partners

**DOI:** 10.1101/2025.07.23.666280

**Authors:** Fjoralba Zeqiri, Thomas Schmidt, Yuko Misumi, Yohei Miyanoiri, Nicole Frankenberg-Dinkel, Genji Kurisu, Eckhard Hofmann

## Abstract

Cyanophages play a critical role in shaping host metabolism during infection by encoding auxiliary metabolic genes (AMGs), including enzymes involved in phycobilin biosynthesis. PebS (phycoerythrobilin synthase) is a phage-encoded ferredoxin-dependent biliverdin reductase (FDBR) that catalyses the two-step reduction of biliverdin IXα (BV) to phycoerythrobilin (PEB). Although high-resolution structures are available for both PebS and its electron donor ferredoxin (Fd), the structural details governing their interaction remain unclear. This study leverages the well-characterised structural models of PebS and a host-like ferredoxin from *Thermosynechococcus elongatus* (Te-Fd), whose NMR structure provides a reliable basis for probing protein– protein interactions. Using [¹⁵N]-labelled, gallium-substituted Te-Fd, we employed NMR spectroscopy to monitor chemical shift perturbations upon binding, enabling us to probe the interaction interface in solution. Based on these data, we conducted protein-protein docking with HADDOCK to predict the interaction interface between Te-Fd and PebS. PebS variants designed to disrupt this interface did indeed show corresponding alterations in enzymatic efficiency and product formation as determined by time resolved UV/Vis spectroscopy and HPLC analyses. Utilizing the Fd encoded in the cyanophage PSSM2-Fd in our assays, we could observe a significantly improved catalytic activity, suggesting an coevolution of phage enzyme and electron donor. A comparison of the two available X-ray structures of Te-Fd and PSSM2-Fd with an alphafold-model of the Fd of the natural host *Prochlorococcus* NATL1A (NATL1A-Fd) also supports this evolutionary adaptation and the role of both PSSM2-Fd and PebS as AMGs involved in viral infection by PSSM2.

## Introduction

Marine cyanobacteria are among the most abundant photosynthetic organisms on Earth.^1^ Their photosynthetic activity makes them central contributors to global primary production and is central to sustaining the carbon and nitrogen cycles throughout the ocean.^2,3^ The productivity of these organisms is tightly dependent on light availability, which drives the evolution of the photosynthetic apparatus and the biosynthetic pathway for pigments. ^4–9^ Cyanophages coevolve to exploit these light-dependent adaptations within a broader viral strategy involving auxiliary metabolic genes (AMGs).^10–14^ These host-like genes are temporarily expressed during infection to maintain or redirect metabolic pathways that are essential for viral replication. ^13,15^ Among common cyanophage-encoded AMGs are those involved in the biosynthesis of bilin-derived chromophores, such as phycoerythrobilin (PEB).^13,15–18^ These linear tetrapyrroles serve as light-harvesting cofactors in phycobiliproteins, which are key components of the photosynthetic antenna in cyanobacteria.^19–21^ In host organisms, PEB is synthesised from the reduction of biliverdin IXa (BV) via a two-step enzymatic reaction involving PebA and PebB, two ferredoxin-dependent biliverdin reductases (FDBRs).^22^ First, PebA reduces BV to 15,16-dihydrobiliverdin (15,16-DHBV), followed by PebB, which completes the reduction of 15,16-DHBV to PEB (Figure 1).^22,23^ In the cyanophage *Prochlorococcus* PSSM2, the two-enzyme system is replaced by PebS, which catalyses the two times two-electron reduction of BV to PEB in a single step, functionally bypassing the two-enzyme route via PebA and PebB (Figure 1).^24–27^ This functional efficiency likely exemplifies the evolutionary pressure to adapt metabolic processes and allow the cyanophage to utilise host resources during its brief replication cycle.^25,26,28^ For PebS to function, it requires ferredoxin (Fd) as an electron donor, analogous to the host-derived FDBRs (PebA and PebB) (Fig. 1). The phage PSSM2 encodes its own Fds, implying a potentially specialised phage-adapted Fd system. Notably, no structural data is available for any FDBR-Fd complex.^29,30^ This lack of atomic details limits our understanding of the molecular details of recognition and binding of PebS’s redox partner. To structurally investigate the interaction site, we leveraged the availability of high-resolution structures for PebS and a host protein like Fd derived from *Thermosynechococcus elongatus* (Te-Fd). The Te-Fd was selected as a well-characterised host-like Fd due to its high similarity with the cyanophage-derived Fd (PSSM2-Fd), well-characterised nature as determined by NMR studies, and its prior use in defining protein-protein interaction sites.^31–38^ These made the Te-Fd a valuable model for studying the interaction with PebS beyond its native viral context. We used NMR spectroscopy to explore the interaction between PebS and Te-Fd.^37^ Building on structural interaction data, we introduced point mutations on PebS which were predicted to mediate binding with Te-Fd. We tested the catalytic activity of the wild-type PebS (wt-PebS) and variants using both the Te-Fd and PSSM2-Fd as electron donors. This dual approach enabled us to evaluate the PebS-Fd interaction site and assess its adaptability to different redox partners. These results provide an advance in the understanding of AMG-driven adaptations and are also used to determine whether PebS exhibits enhanced catalytic performance with its native PSSM2-Fd, suggesting possible viral adaptation. In addition, we considered the native host ferredoxin from *Prochlorococcus* NATL1A (NATL1A-Fd), a member of the same plant-type [2Fe–2S] ferredoxin family. While no experimental structure exists for NATL1A-Fd, a predicted model is included in the structural comparisons conducted in this study.

**Figure 1.**
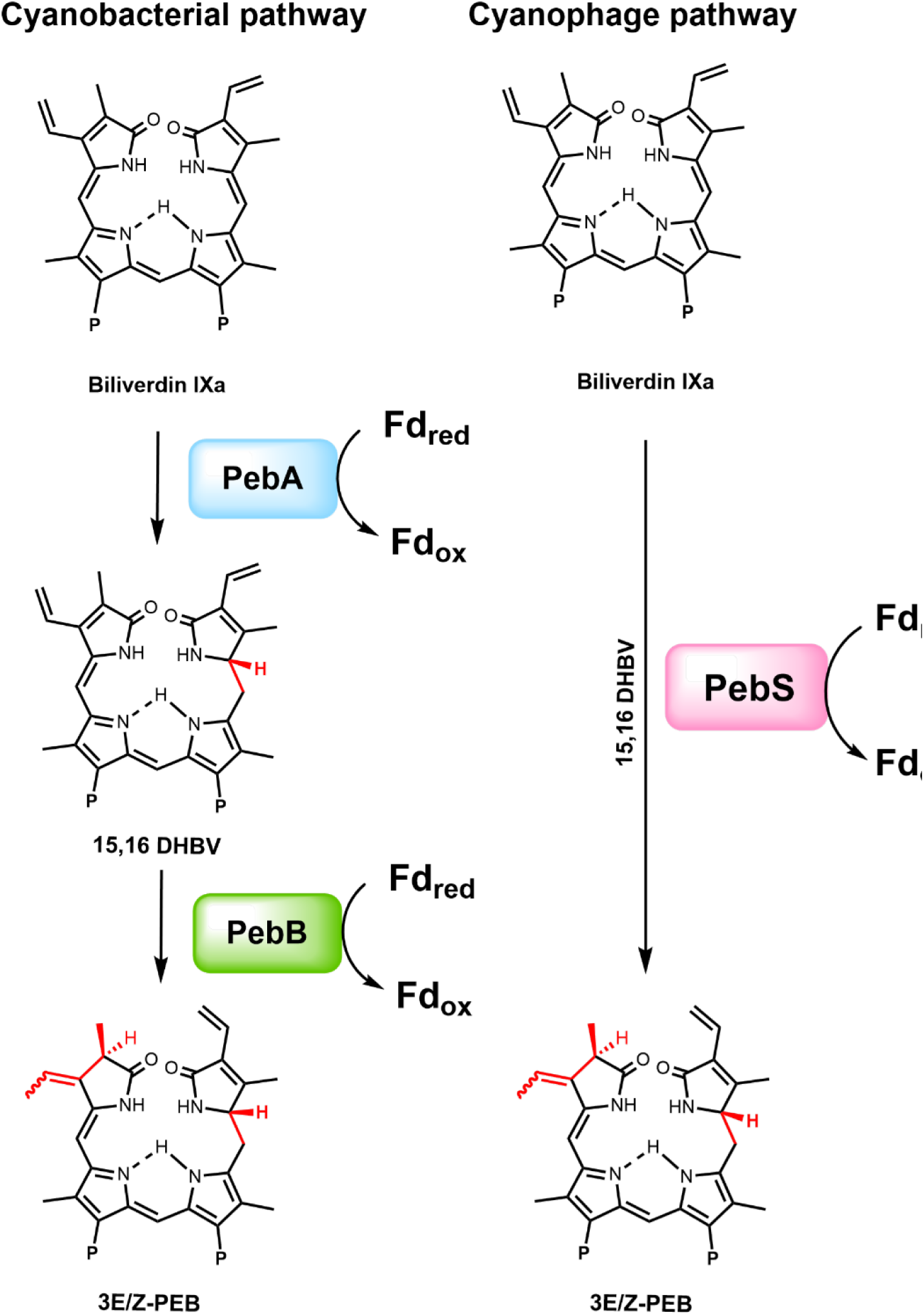
Comparison of canonical (PebA/PebB) and phage-mediated (PebS) PEB biosynthesis pathways. In the host pathway, PebA and PebB sequentially reduce biliverdin IXα (BV) to phycoerythrobilin (PEB) via the intermediate 15,16-dihydrobiliverdin (15,16-DHBV), using ferredoxin (Fd) as the electron donor. In contrast, phage-encoded PebS catalyzes both steps in a single reaction, producing the same intermediate and final product with Fd as the electron donor. The (P) abbreviation stands for propinoate groups and the Fdox and Fdred stands for oxidised and reduced ferredoxin.

## Material and methods

### Reagents

All chemicals were bought from Sigma Aldrich unless otherwise specified.

### Mutagenesis, expression and purification of PebS, variants, and Fds

The mutagenesis for PebS variants was done using the NEBaseChanger assist tool (Table S1). The PCR reactions were then completed using the Q5 High-Fidelity 2X master mix. The pGEX-6P-3 vector system was used for cloning and expressing PebS and its variants. Expression and purification of GST-tagged PebS and variants were performed as described earlier, with some minor adjustments.^28,39^ Protein expression was performed in TB medium until the OD_600_ reached 0.8. The culture was placed in an ice bath for 1 hour before induction with 500 µM IPTG, then it was left to shake overnight at 16 °C. The protein expression was interrupted by harvesting with a SLC-6000 rotor at 4000 g for 10 minutes at 4 °C. The pellets were stored at −20 °C. For protein isolation, the cells were resuspended in lysis buffer (50 mM Tris/HCl, pH 8, 300 mM NaCl) at a ratio of 1:10 per gram of cell. In the lysis buffer, prior cell disruption 1 mM PMSF was added. The cell lysate was homogenised using a douncer, and cell disruption was carried out by 3-5 cycles through a LM10 Microfluidizer® High Shear fluid homogenizer (Microfluidics) at 800-1000 bar. The resulting mixture was subjected to ultracentrifugation in a Ti45 rotor at 10000 g at 4 °C on an Optima L-80 XP centrifuge. The supernatant was applied to a Glutathione Sepharose (GSH) column (GSTrap HP 5 ml) column on an Äkta purifier system (GE Healthcare). Elution was performed using lysis buffer containing 25 mM reduced glutathione. Protein-containing fractions were combined followed by addition of precision protease and were dialysed in lysis buffer overnight. Uncleaved protein and protease were removed by a second GSH step. The flowthrough was collected and concentrated to 1-2 mM for size exclusion chromatography (HiLoad 16/60 Superdex 75 Prep Grade, GE Healthcare). Peak fractions were pooled and concentrated to 1-2 mM, then frozen for further use. Fd and [N^15^] gallium substituted Fd were purified as described earlier.^37^ The protein sample was concentrated to 1 mM and frozen for further use.

### NMR measurement and docking

Fd-Pebs and Fd-PebS-BV were measured in 20 mM Na_3_PO_4_, pH 6.8, and 50 mM NaCl, as described earlier.^37^ NMR measurements were conducted at 293K on an Avance III HD 800 spectrometer (^1^H resonance frequency: 800 MHz; Burker, Billerica, MA, USA) equipped with a TXI cryogenic probe. The resonance assignments of the [^15^N] Fd, prepared at a final concentration of 300 µM, were carried out as previously described.^37^ Titration experiments with PebS were also perform, as described in a previous report.^37^ The optimal measurement condition was determined to be a 1:2 molar ratio of [^15^N] Fd to PebS. Subsequently, BV was added at the same ratio as PebS. NMR data were processed using Topspin 3.6.2 (Bruker) and NMRFAM-Sparky.^40^ Chemical shift perturbations (Δδ) of [^15^N] Fd upon adding of PebS and BV were calculated using the following equation: Δδ = [(Δδ1 HN) 2 + (0.04Δδ15 N) 2 ] 1/2 where δ1 HN and δ15 N represent the chemical shifts of amide proton and nitrogen for each amino acid residues, respectively (Fig. S4).^37^ Docking simulations were performed using HADDOCK, based on previously published structures of Fd and PebS (PDB ID 5AUI and 2VCK, respectively).^28,37,41^ Protein-protein interfaces were identified from the HADDOCK docking models using LigPlot+, which provided structural insights to guide the design of PebS point mutants (Fig. S5).^42^

### Anaerobic reductase assay and HPLC measurements

The UV/Vis spectroscopic measurements of the anaerobic reductase assay were performed as already described with minor adjustments.^24^ As electron donor the host-like Fd (Te-Fd), which was utilised during the NMR studies, and the phage-derived ferredoxin PSSM2-Fd were used. Reactions were quenched after 20 min using 0.1% trifluoracetic acid (TFA). The resulting products were isolated using C-18 Sep-Pak (Waters) as per manufacturer’s instructions and lyophilized. HPLC measurements were carried as described earlier.^24,43^

## Results and discussion

### Identification of the [^15^N] Ga-Ferredoxin-PebS interaction sites and in silico-docking

Using NMR spectroscopy, we identified perturbations of host-like ferredoxin Te-Fd resonances upon titration of PebS in the presence and absence of the substrate BV, as depicted in (Fig. 2, Fig. S2. Fig. S3). Notably, residues C40, R41, S46 and D66 exhibited significant chemical shift changes upon addition of PebS, indicating a strong interaction with the enzyme (Fig. S4). In addition to these key residues, moderate chemical shift changes were observed for A42, D61, S60, and Q69 ,suggesting their involvement in an interaction that may contribute to the stabilisation of the binding interface. Upon BV addition, the key residue profile remained consistent with C40, R41, and D66 exhibiting the dominant shifts, which indicated that the interaction epitope remained essentially unchanged (Fig. 2, Fig. S4). Interestingly, L65, which showed no perturbation in the absence of BV, demonstrated a moderate change in the presence of the substrate, suggesting its specific involvement in the BV-bound state (Fig. 2). In contrast, S46, which initially showed strong perturbation, no longer appeared to participate in the interaction under BV-bound conditions. Furthermore, minor chemical shift changes were observed for E21, D22, E23, Q32,A42, D61, S60, Q62, and Q69 upon BV addition, implying that these residues may enhance the overall binding affinity between PebS and Te-Fd.

**Figure 2.**
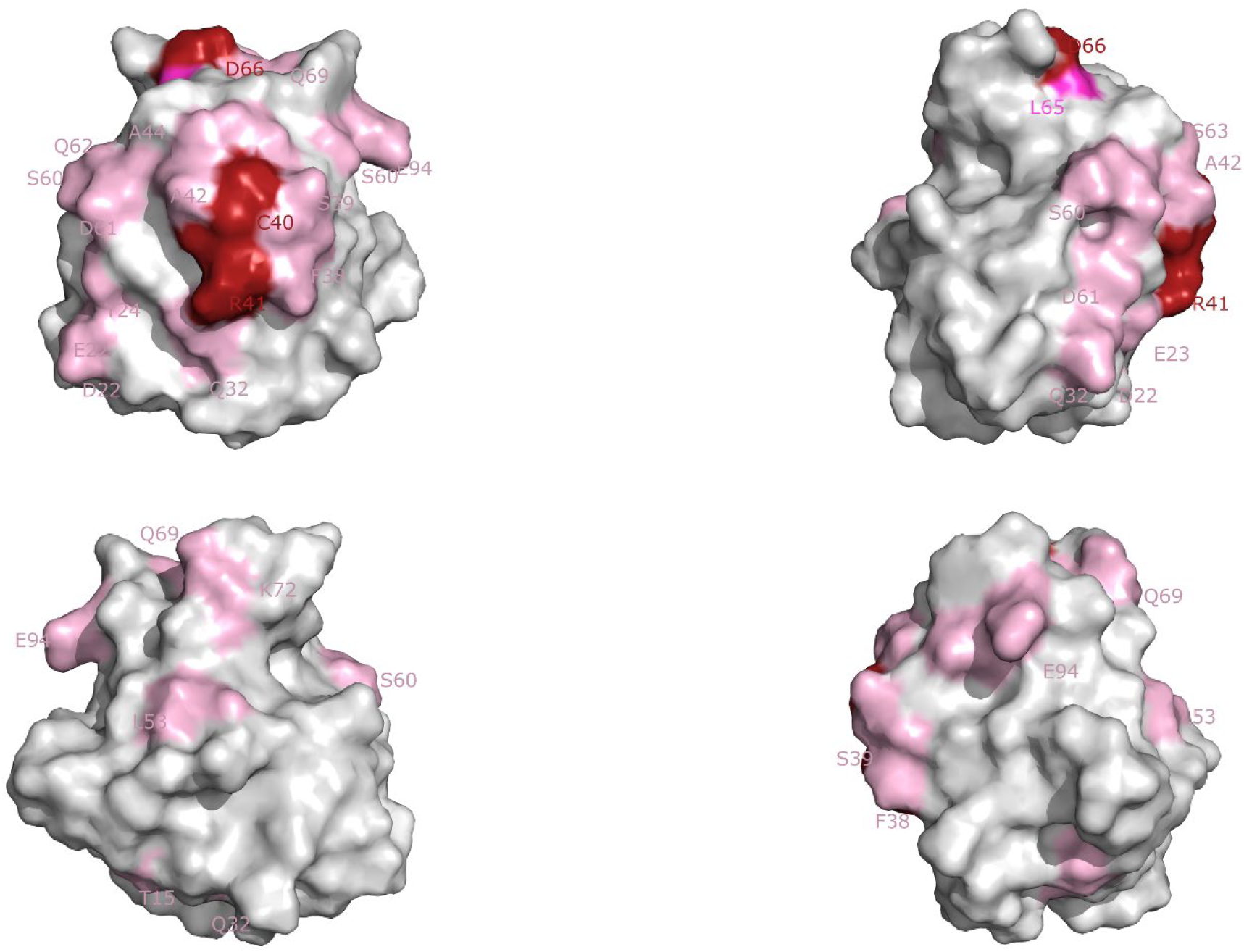
Observation of chemical shift perturbation of [^15^N] Ga replaced Te-Fd. Chemical shift changes upon PebS and BV addition. Interaction regions with PebS include C40, R41, L65, D66. Shown is the accessible surface for Te-Fd in 4 different orientations, rotated by 90° around the vertical axis. Colour mapping of the residues: light pink 0.05-0.01 ppm; pink 0.01-0.015 and red above 0.015.

Importantly, residues C40, R41, and D66 consistently exhibited pronounced chemical shift changes under both conditions, strongly supporting their direct involvement in the interaction with PebS (Fig. 2, Fig. S3, Fig. S4).

These data indicate that the central binding epitope on Te-Fd for the interaction with PebS is largely maintained in the presence of substrate, with only minor modulations of the local binding environment. The 2Fe2S-cluster is located right in the center of this epitope, thereby reducing the distance for electron transfer towards PebS (Fig. S11). Very similar interaction epitopes have been observed for other plant type Fds in the interaction with e.g. photosystem I, plant heme oxygenase, nitrite reductase or ferredoxin-NADP+ reductase. ^31,34–36,38,44–46^

### Structure-guided selection of point mutations on PebS based on protein-protein docking

To identify the corresponding PebS interaction epitope, we then docked PebS to Te-Fd whilst defining C40, R41, L65 and D66 as active restraints in HADDOCK docking simulations.^41^ We analysed the resulting model with LigPlot+ to identify pairs of residues on both proteins predicted to contribute to the interaction (Fig. S5).^42^ Our docking results identified eleven surface-exposed residues likely contributing to binding to Fd.

From these, we selected eight residues for site-directed mutagenesis: F109Y, K113E, F143Y, L205E, G210A, Y211F, K213E,and N214D, and. The variants were designed to subtly or significantly alter the interface properties, such as hydrophobicity, charge, or hydrogen bonding potential. We excluded residues previously implicated in binding to the BV cofactor (N83, P207, R209) to maintain the enzyme’s catalytic functionality.^24,28,47–49^ All PebS variants showed expression levels, elution profiles and solubility during purification comparable to the Wild-type PebS (wt-PebS).

### Anaerobic reductase assays with host-like ferredoxin, Te-Fd as electron donor

Purified PebS and variants were characterised with the anaerobic reductase assay as previously described.^24^ First we utilized the host protein like ferredoxin Te-Fd as the electron donor, which we had used previously in our NMR structural studies. In this setup, Wt-PebS efficiently catalysed the conversion of BV to phycoerythrobilin (PEB), as confirmed by the appearance of characteristic absorbance peaks in the UV/Vis spectra and HPLC analysis of the reaction products (Fig. S7A). Variants F143Y, G210A and Y211F showed substrate conversion levels comparable with wt-PebS (Fig. S8A-S10A). Variant L205E exhibited slightly slower conversion of BV, as demonstrated by the reduced absorbance intensity and lower peak area in the HPLC chromatograms (Fig. 3).

**Figure 3.**
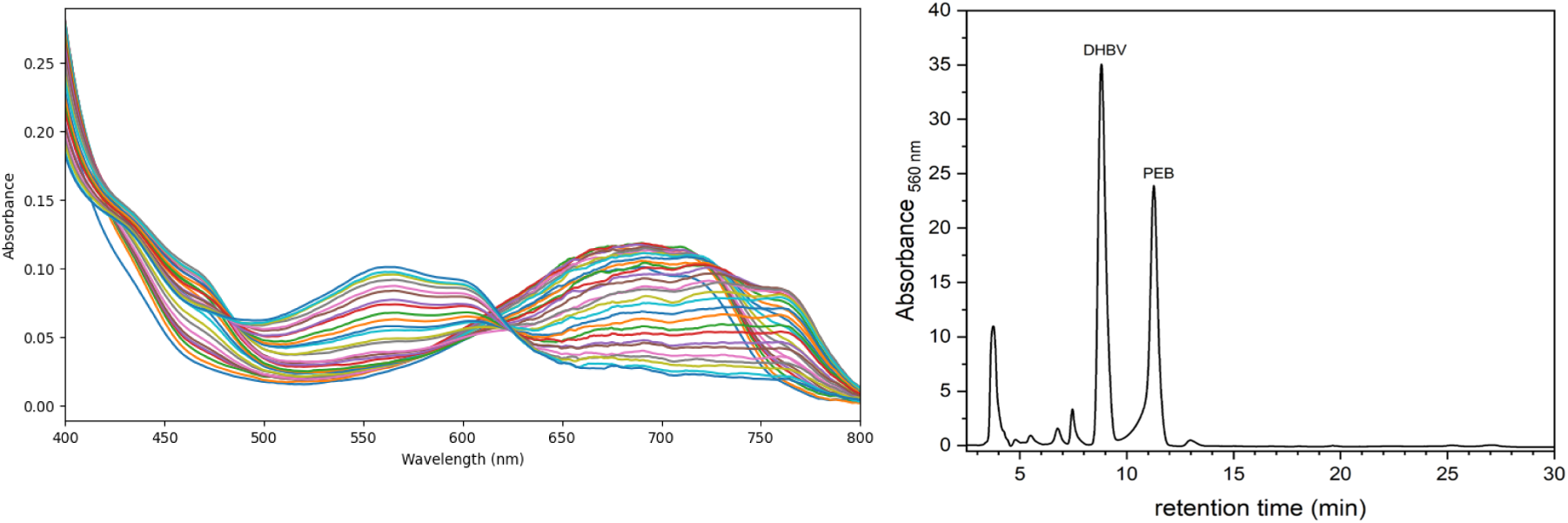
Spectroscopic and chromatographic analysis of the PebS L205E variant with Te-Fd. UV/Vis absorbance spectra of the enzymatic reaction. Phycoerythrobilin (PEB) shows a characteristic absorption peak near 550 nm, while unconverted biliverdin IXα (BV) absorbs around 650 nm, consistent with values reported by Dammeyer et al., 2008. HPLC chromatogram of the quenched reaction mixture, confirming product formation and separation between PEB and residual BV.

In contrast, variants K113E, K213E and N214D demonstrated markedly reduced BV conversion efficiency (Fig. 4). These variants accumulated the intermediate DHBV with only low PEB formation.

**Figure 4.**
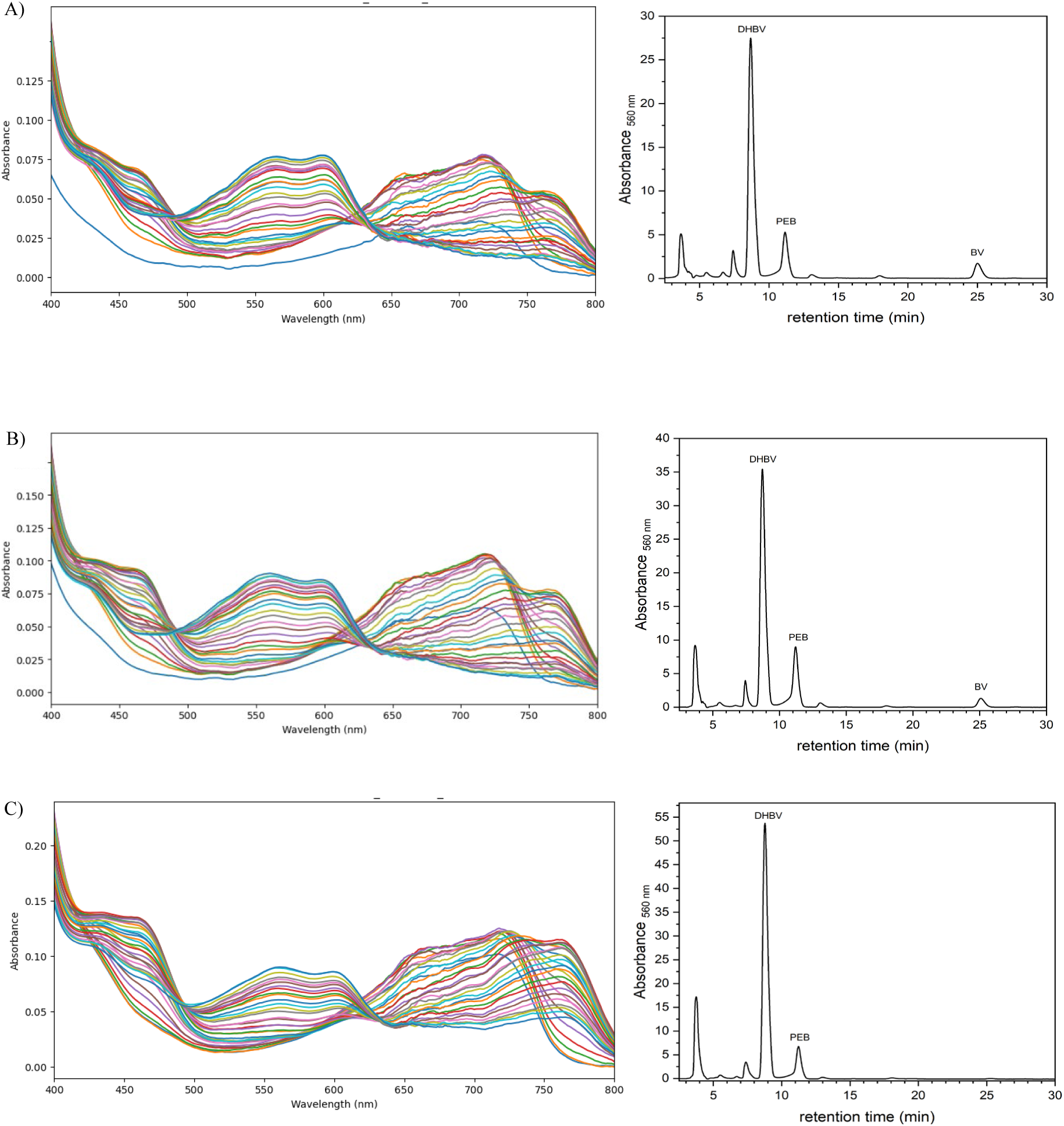
Spectroscopic and chromatographic analysis of PebS variants K113E, K213E, and N214D with Te-Fd. Each panel presents a UV/Vis absorbance spectrum (left) and corresponding HPLC chromatogram (right) of the enzymatic reaction. A) K113E; B) K213E; C) N214D; In all UV/Vis spectra, phycoerythrobilin (PEB) exhibits a characteristic absorption peak near 550 nm, while biliverdin IXα (BV) absorbs around 650 nm, consistent with Dammeyer et al., 2008. HPLC traces confirm product formation and show separation of reaction components, with peaks corresponding to 15,16-dihydrobiliverdin (DHBV), PEB, and residual BV.

This data confirm that the predicted salt bridges and polar interactions these residues are involved in (Fig. S5) are essential contributors for interface stability.

Variant F109Y exhibited complete loss of activity with no detectable PEB formation (Fig. 5). This dead variant suggests that the F109 residue plays a critical role in completing the interaction epitope between substrate-bound PebS and Fd. The addition of a hydroxyl group seems to interfere with efficient complex formation.

**Figure 5.**
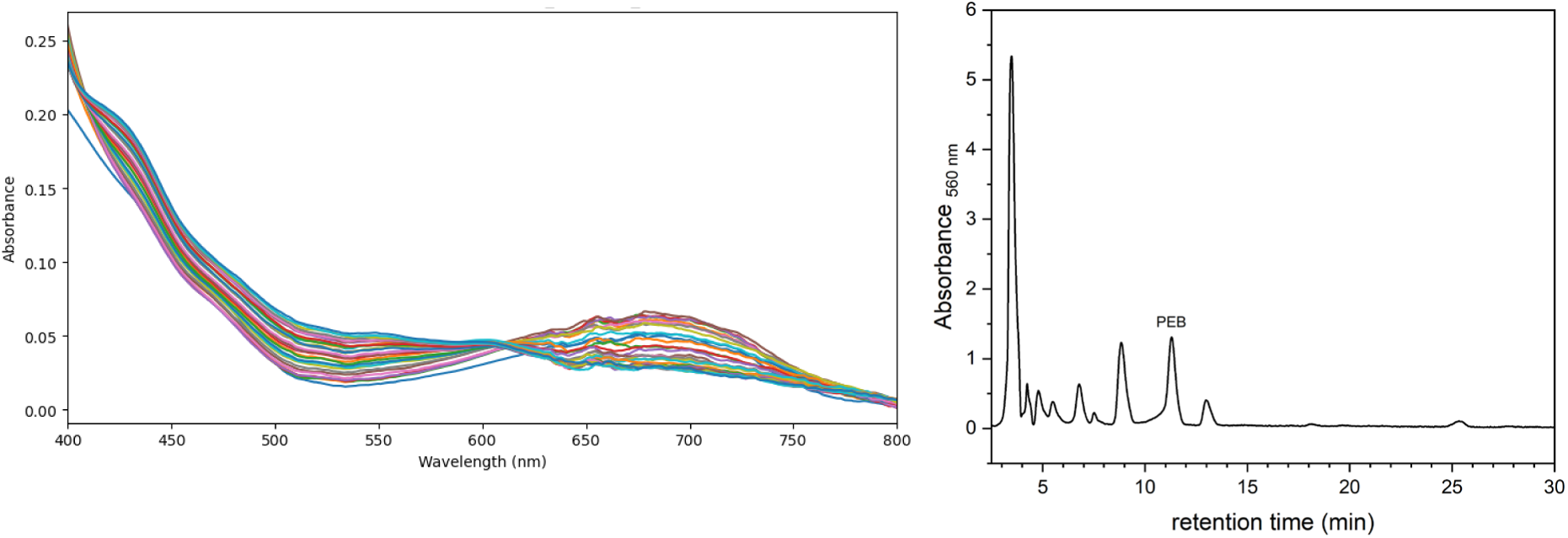
Spectroscopic and chromatographic analysis of the PebS F109Y variant with Te-Fd. The panel presents a UV/Vis absorbance spectrum (left) and corresponding HPLC chromatogram (right) of the enzymatic reaction. UV/Vis and HPLC data indicate no detectable enzymatic activity under the tested conditions. No product peaks corresponding to 15,16-dihydrobiliverdin (DHBV) or phycoerythrobilin (PEB) were observed.

The experimental results support a broader trend: minor side chain modifications tend to preserve activity, whereas mutations at key residues involved in salt bridge formation or major hydrophobic interaction patches often result in severe loss of activity or complete functional loss. This highlights the importance of electrostatic and hydrophobic interactions for the proper formation of the PebS-Fd complex and further support the proposed interaction model.

### Anaerobic assays with PSSM2-Fd as electron donor

As already described, for NMR studies, we had to use the Te-Fd as an established system; however, in the biological context, PebS will likely interact with the phage-encoded Fd (PSSM2-Fd). We therefore repeated our activity assays, now using PSSM2-Fd as an electron donor. Interestingly, we observed significantly improved catalytic rates for all PebS variants except F109Y. The improvement in the kinetics was observable in both the UV/Vis and the HPLC results, revealing an elevated PEB peak (Fig. 6). Variants K113E, K213E and N214D even partially regained the ability to reduce BV to PEB entirely. Since the PSSM2-Fd and PebS are co-expressed within the same host during phage infection, these two proteins have likely coevolved to form an optimised and more efficient electron transfer.^24^ This is in line with earlier findings, where PebS showed significantly increased catalytic turnover with PSSM2-Fd when compared to another cyanobacterial Fd from *Synechococcus* sp. P7002.^24^

**Figure 6.**
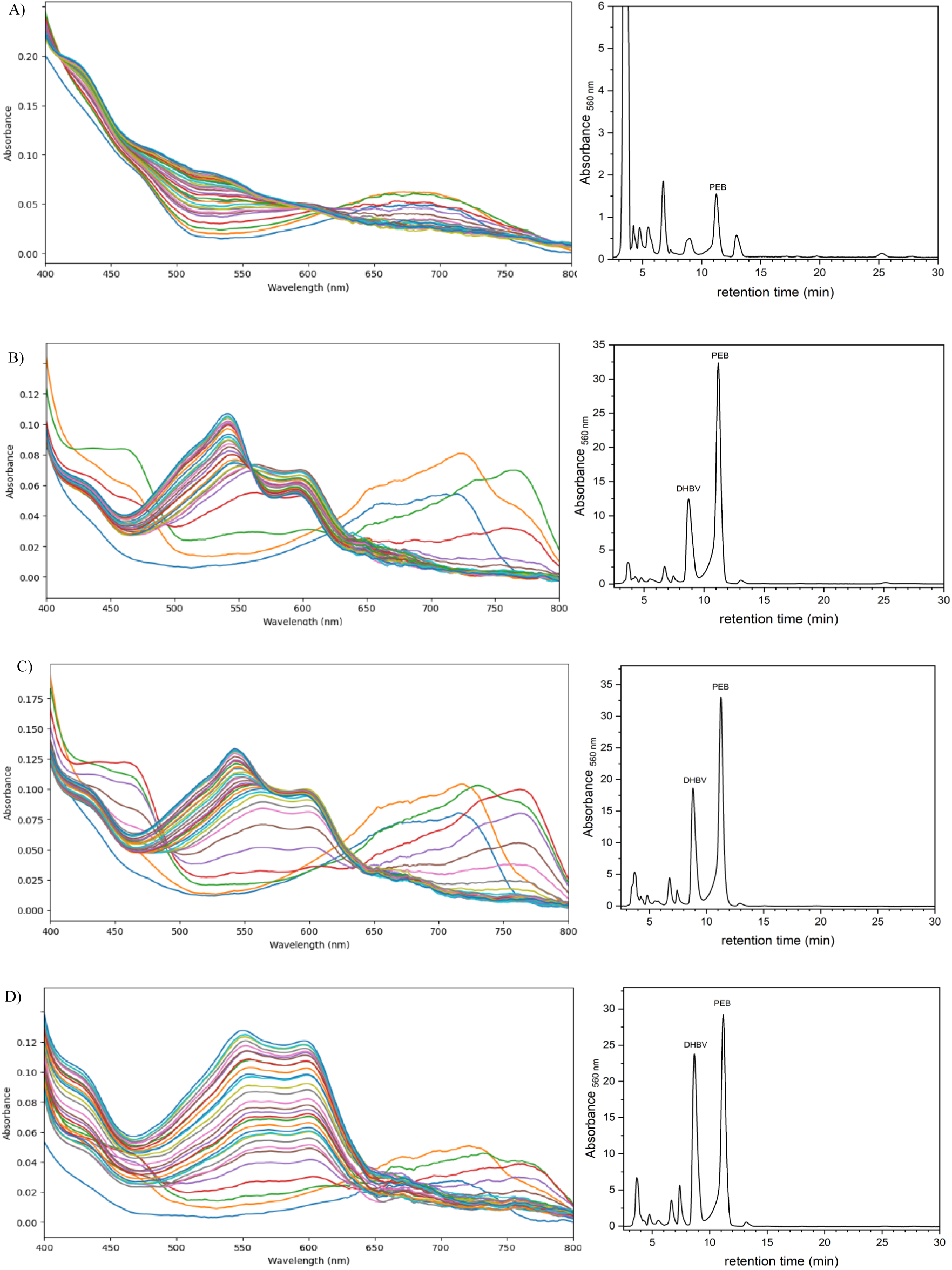
Spectroscopic and HPLC analysis of PebS variants with native PSSM2-Fd. UV/Vis absorbance spectrum (left) and corresponding HPLC chromatogram (right) of different variants: A) F109Y B)K113E C) K213E D) N214D. In each panel, absorbance peaks indicate pigment formation, with phycoerythrobilin (PEB) absorbing near 550 nm and biliverdin IXα (BV) around 650 nm, as reported by Dammeyer et al., 2008. HPLC traces show elution profiles of reaction products; DHBV and PEB peaks are marked accordingly

### Comparison of Ferredoxins from Te-FD, P-SSM2-FD, and *Prochlorococcus* NATL1A

To better understand the observed differences in interaction efficiency, we compared the two ferredoxins and also included the ferredoxin from the native host *Prochlorococcus* NATL1A. All three sequences share a high degree of conservation for core residues, specifically of the cysteine residues contributing to the [2Fe-2S] cluster binding (e.g., C40, C45, C48, C78 (Te-Fd numbering) (Fig. 7). The key residues identified in the NMR titration experiments on Te-Fd (C40, R41, D66, S46, L95) are conserved in PSSM2-Fd and are also at least conservatively replaced in NATL1A-Fd. NATL1A-Fd features two distinct insertions outside the core that are not present in either Te-Fd or PSSM2-Fd. While Te-Fd and PSSM2-Fd show a very high overall sequence identity of 60%, NATL1A-Fd has diverged stronger, as we observe only 38% and 28% identity to TeFd and PSSM2-Fd, respectively (omitting the two sequence insertions in the calculation).

**Figure 7.**
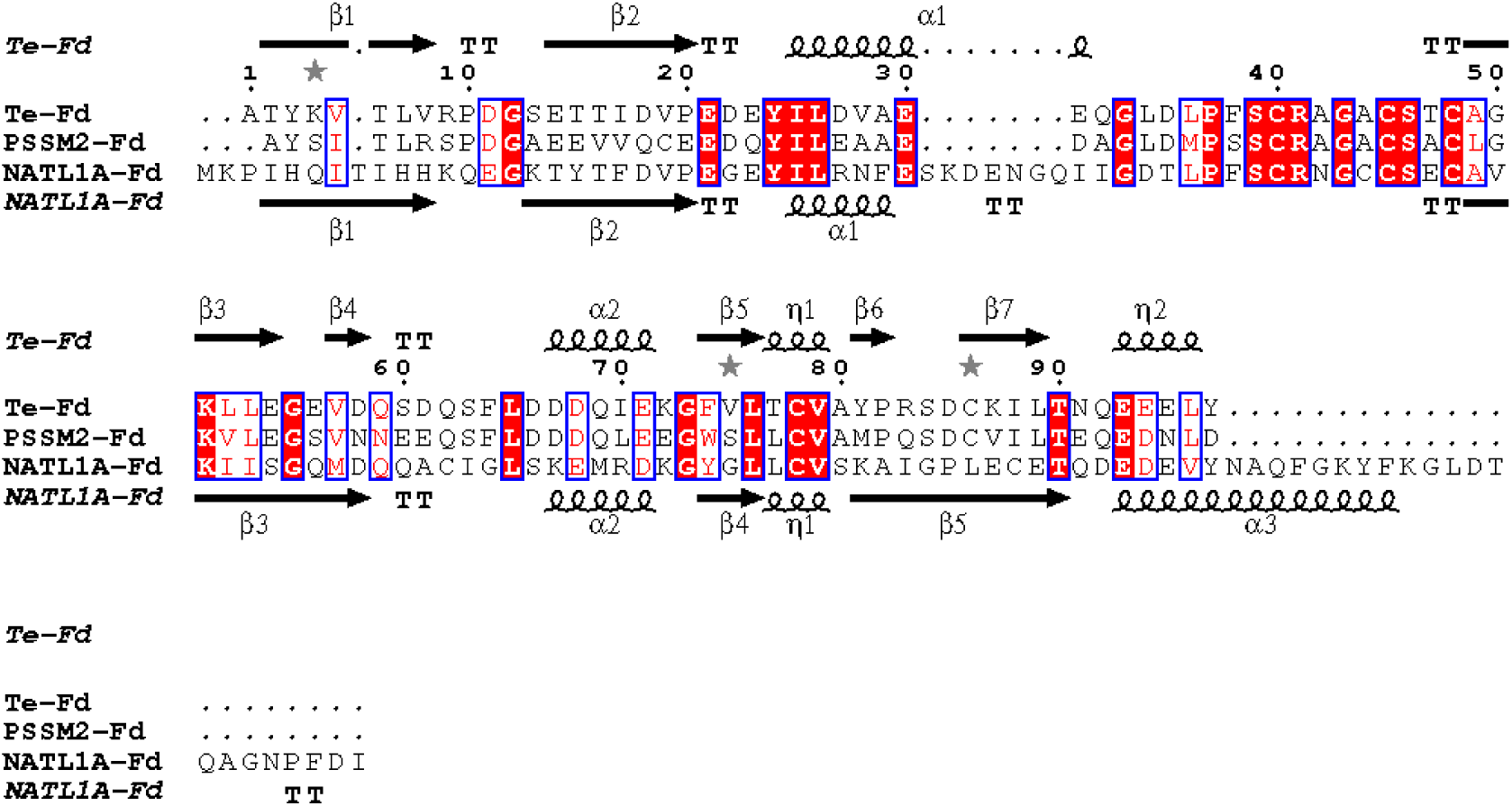
Amino acid sequence alignment of Te-Fd, PSSM2-Fd and NATL1A-Fd. Conserved residues are shaded in red. Secondary structure elements of Te-Fd (PDB-ID 5AUI) and of NATL1A-Fd (alphafold2-model) are shown above and below the alignment. Figure produced with ESPript.^50^

We also compared the 3D-structures of Te-Fd (PDB ID: 5AUI), PSSM2-Fd (PDB ID: 6VJV) and an Alphafold-Model of NATL1A-Fd (Fig. S11).^51^ All three ferredoxins share the highly conserved core fold characteristic of plant-type [2Fe–2S]. Consistent with the high sequence similarity, Te-Fd and PSSM2-Fd also show nearly identical backbone conformation (Root mean square deviation (RMSD) of 0.45 Å for 84 aligned Cα atoms)(Fig SX). NATL1A-Fd still superposes fairly well in the core backbone region (RMSD 0.77 Å to for 75 aligned Cα atoms and 0.88 Å for 76 aligned Cα atoms to Te-Fd and PSSM2-Fd, respectively). On the all atom level, Te-Fd and PSSM2-FD also superpose very well, but NATL1A-Fd clearly diverged more, due to the lower sequence conservation and the two insertions.

Commonly the highly charged surface of Fds is discussed to be the major contributor to interaction specificity in Fd dependent systems. We therefore calculated the APBS electrostatic potential map for all three proteins (Fig. 8).

**Figure 8.**
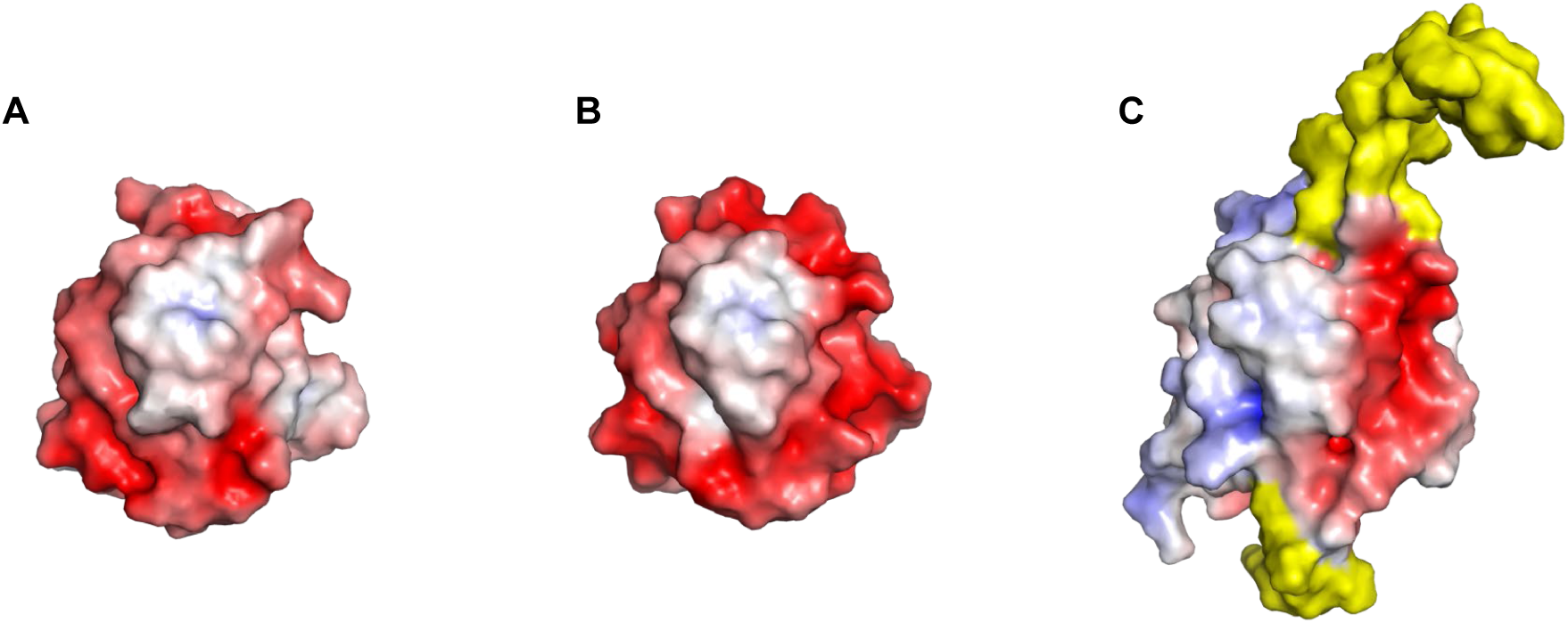
Comparison of ferredoxin surface polarity around the interaction epitope. Electrostatic surface potentials generated using APBS and visualised in PyMOL for Te-Fd (A), PSSM2-Fd (B) and NATL1A-Fd (C). Orientation analogous to figure 8. Red indicates negative potential, blue indicates positive potential, and white indicates neutral regions. Sequence insertions in NATL1A-Fd shown in yellow. While Te-Fd and PSSM2-Fd display highly conserved surface charge distributions, especially in regions associated with functional interfaces, NATL1A-Fd shows larger modifications both in shape and polarity, despite the overall highly conserved fold (see Figure S:11).

Both Te-Fd and PSSM2-Fd exhibit similar electrostatic profiles and share conserved residues at their predicted binding interfaces. However, we observed a significant difference in their ability to support the catalytic activity of PebS. Specifically, the use of PSSM2-Fd in the activity assays produced higher levels of product formation across both the wild-type and several mutant variants of PebS. The data imply that successful redox interactions require more than charge matching or conserved contacts, but they depend on precise interface geometry and optimal alignment of the redox sites

One possible explanation for the superior performance of PSSM2-Fd is its co-evolution with the native PebS, resulting in a redox partnership that is optimised both structurally and functionally. This optimisation may encompass better alignment of redox-active centres and improved surface complementarity, facilitating faster or more efficient electron transfer. In comparison, our data do not directly confirm co-evolutionary adaptation; PSSM2-Fd’s ability to partially restore activity in impaired PebS variants, where Te-Fd does not, provides compelling functional evidence that such adaptation is biologically significant.

In the physiological situation of an phage infection, PebS would also have the option to interact with the host Fd. Both structural and sequence data clearly indicate, that NATL1A-Fd diverged strongly from both Te-Fd and PSSM2-Fd. This is very prominent in the polar surface representation in Fig. 8, where clear deviations in both surface polarity and shape can be observed as compared to the other two ferredoxin.

Yet, NATL1A-Fd more closely resembles Te-Fd than PSSM2-Fd, especially in the region modeled for PebS contact, which shows higher variation in PSSM2-Fd. A possible contribution seems less central in docking models (data not shown). Thus, although we cannot entirely dismiss the effect of the NATL1A-specific insertions on PebS binding, the available data imply that the phage enzyme and its corresponding ferredoxin have co-optimized their functions. Based on sequence identity, conserved contact features and structural similarity, NATL1A-Fd is more similar to Te-Fd than to PSSM2-Fd. We therefore expect that PebS will show lower activity also with the native host ferredoxin.

In summary, our data suggest a coevolution of PebS and PSSM2-Fd as auxiliary metabolic genes. Given that the phage host encodes a significantly divergent ferredoxin, it can also be speculated, that the original gene for PSSM2-Fd has not been derived from *Prochlorococcus*, but rather from a non host cyanobacterium.^52^

## Acknowledgments

We thank Petros Sarantopoulos for excellent technical support. We also like to thank Dr. Federica Frascogna for helpful discussion and technical support. This work was supported by the Deutsche Forschungsgemeinschaft by grants HO2600/3-2 and FR1487/10-2 (to E. H. and N. F.-D.) and funding within the Research Training Group 2341 “Microbial Substrate Conversion (MiCon)” to F.Z. and E.H..

## Author contribution

FZ designed the study, performed protein purifications (including NMR samples), planned biochemical assays, conducted HADDOCK docking and structural visualization, and wrote the manuscript. EH analyzed protein–protein interfaces and edited the manuscript. TS carried out activity assays and HPLC. NFD reviewed and revised the manuscript. NMR measurements were performed by YoM. The FD purification protocol was based on methods developed and taught by YuM. NMR experiment preparations were done in GK’s lab, who provided infrastructure and reagents.

