## Supplementary Tables and Figures for "Understanding PebS–Ferredoxin Recognition: A Structural Perspective on Viral and Host Redox Partners"

**Table S1: Primers used for site-directed mutagenesis of PebS.** Each mutation was introduced using a pair of forward (FWD) and reverse (REV) primers targeting specific amino acid residues in the pebS gene. Primer sequences are written 5' to 3'. Where shown, bold letters indicate the codons altered to create the desired mutation.

| Primer | DNA Sequence 5' to 3' |
|---|---|
| K113E_1_FWD | TAGTGATAAG**GAA**GTCATTATAGTATTTG |
| K113E_1_REV | AACTTCATCAGATCCATAC |
| K213E_1_FWD | AGGATATATG**GAA**AATAAG**TTT**GGTG |
| K213E_1_REV | CTAACAGGATCAAGTTCAG |
| L205E_1_FWD | TATGACTGAA**GAA**GATCCTGTTAGAGGATATATG |
| L205E_1_REV | TAAGTATCAAAGTCACTATAGAC |
| N214D_1_FWD | ATATATGAAG**GAT**AAG**TTT**GGTGAG |
| N214D_1_REV | CCTCTAACAGGATCAAG |
| Y211F_1_FWD | TGTTAGAGGA**TTT**ATGAAGAATAAGTTTG |
| Y211F_1_REV | GGATCAAGTTCAGTCATATAAG |
| F143Y_1_FWD | TAAGTATAGA**TAT**TTTGAAATGGGTAATCATTTC |
| F143Y_1_REV | CCATCATCTTCGGGTAG |
| F109Y_1_FWD | TCTGATGAAG**TAT**AGTGATAAGAAG |
| F109Y_1_REV | TCCATACCAAAACAAGG |
| G210A_1_FWD | TCCTGTTAGA**GCG**TATATGAAGAATAAGTTTG |
| G210A_1_REV | TCAAGTTCAGTCATATAAGTATC |

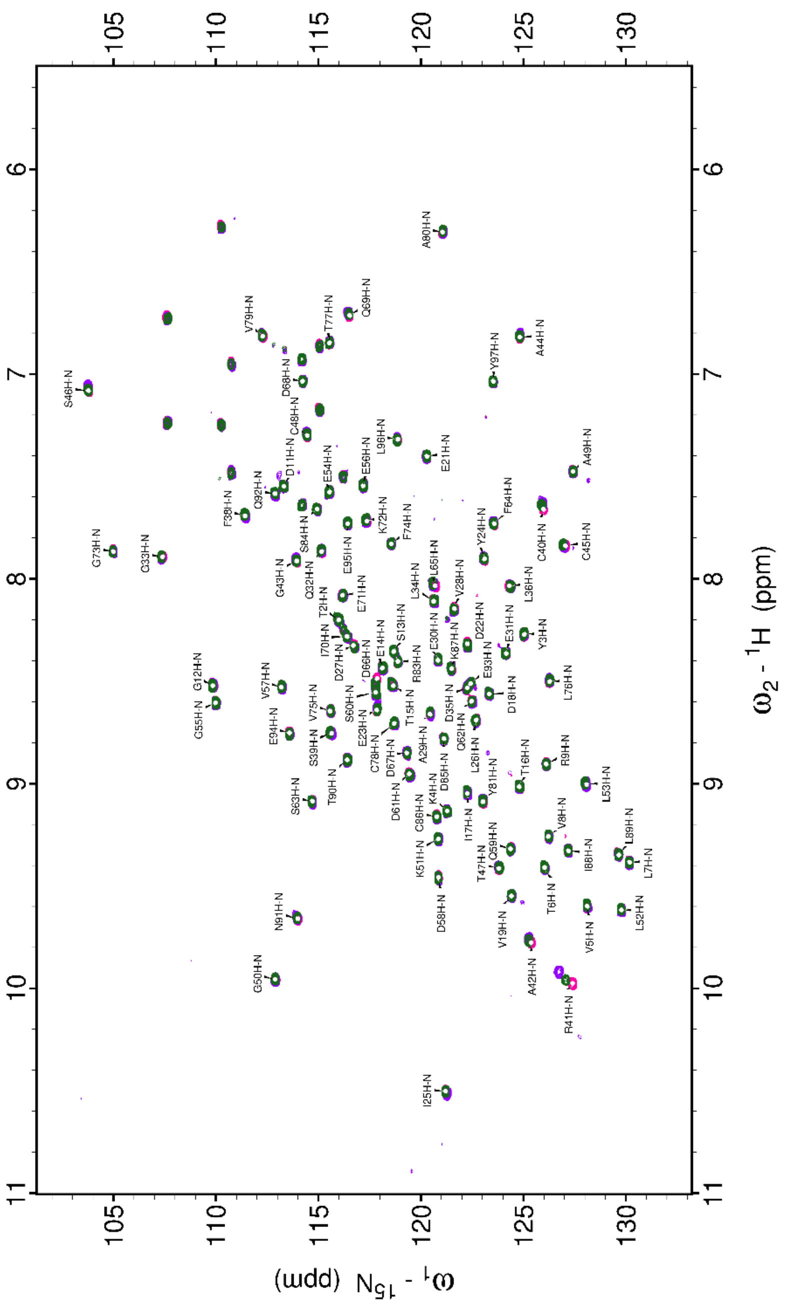


**Figure S2:1H- 15 NSQC spectra of the 15N Ga-Fd.** Overlay of the spectra of free 15 NGaFd (green); in the presence of PebS (purple) and PebS and BV (magenta).

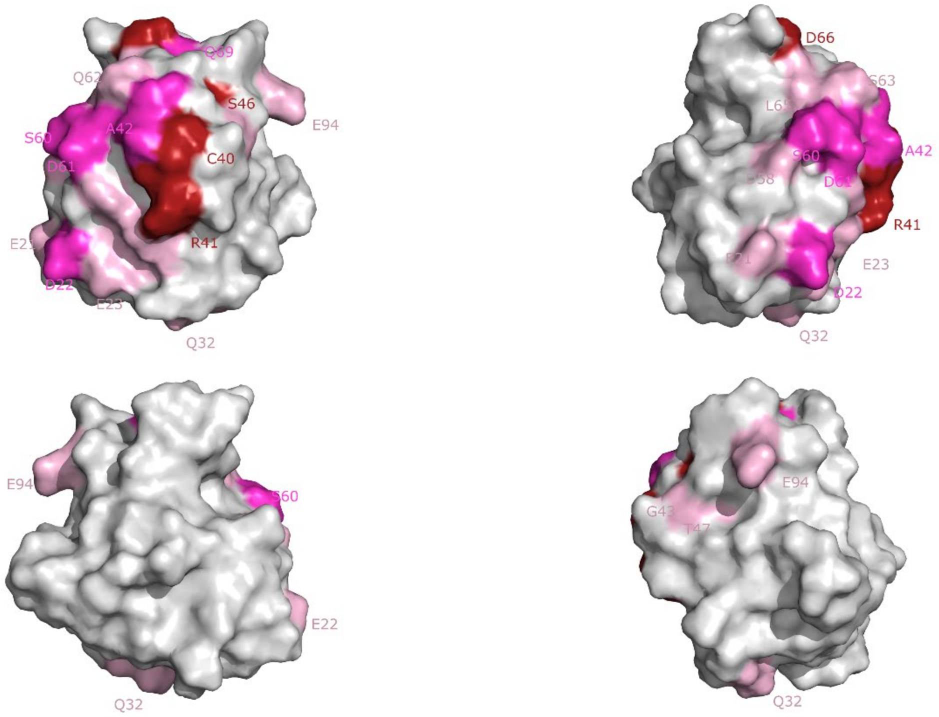


**Figure S3*:*** Mapping of Fd spectral shift intensity during titration with substrate-free PebS. Chemical shift perturbations observed by NMR upon titration of ferredoxin (Fd) with substrate-free PebS are mapped onto the Fd structure. Shift intensity reflects interaction strength across the protein surface, highlighting residues likely involved in PebS binding in the absence of substrate.

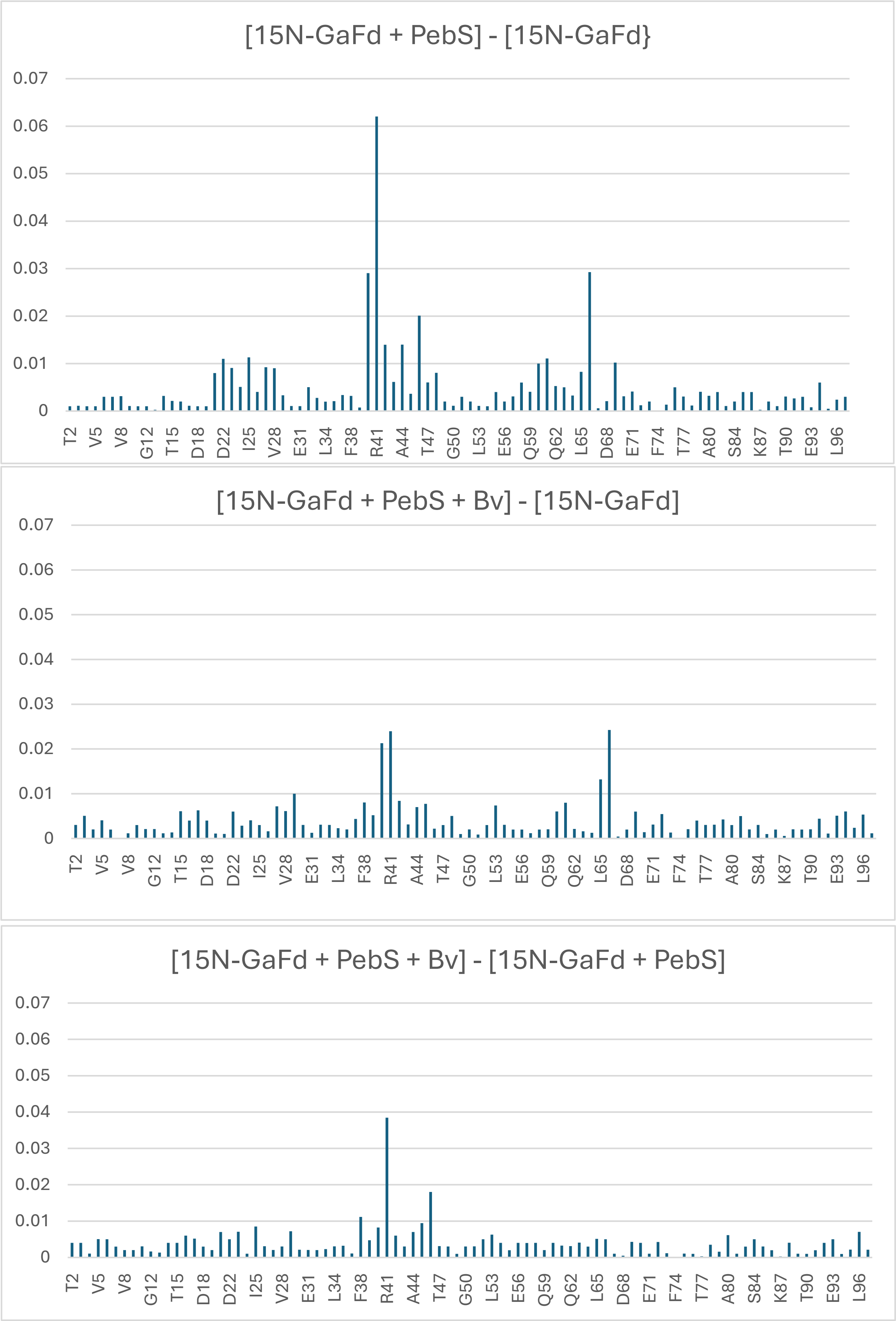


**Figure S4: Chemical shift perturbations between the complex (PebS-GaFd) across various amino acid residues.** The data show residue signal changes of GaFd upon PebS or PebS+BV addition.

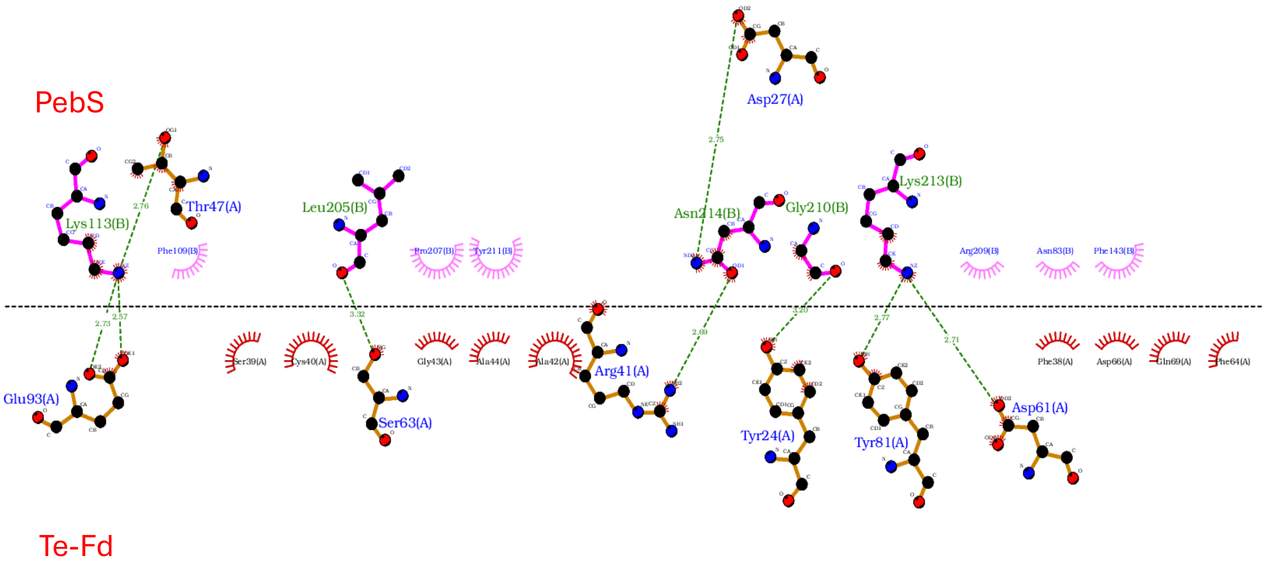


**Figure S5: The visualisation in LIgPlot+ of the PebS and Te-Fd interaction, based on the HADDOCK docking results.**

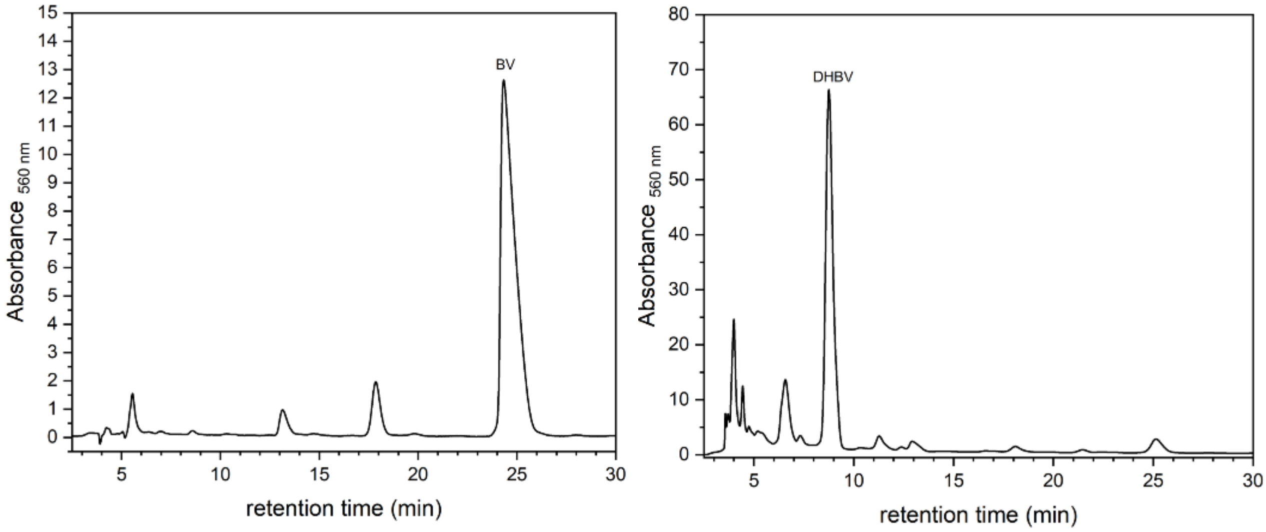


**Figure S6: Reference standards for biliverdin IXα (BV) and 15,16-dihydrobiliverdin (DHBV).** UV/Vis absorbance and HPLC chromatograms of BV and DHBV are shown as standards for comparison with enzymatic reaction products. BV exhibits a characteristic absorbance maximum around 650 nm, while DHBV absorbs slightly blue-shifted, typically around 600 nm. Chromatographic separation was carried out under identical conditions as for the enzymatic assays. Retention times and spectral features are consistent with previously reported values (Dammeyer et al., 2008), confirming the identity and resolution of the bilin intermediates.

### Anaerobic reductase assay

A)

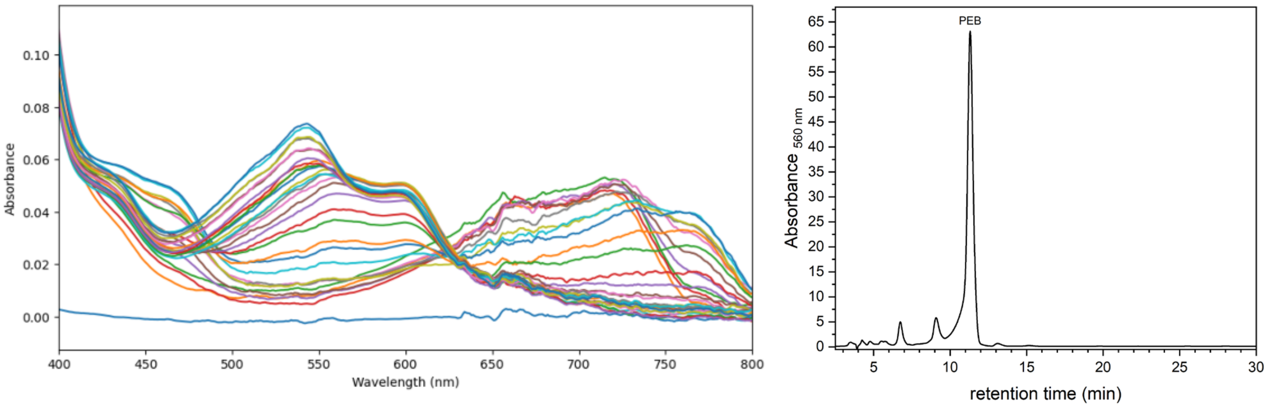


B)

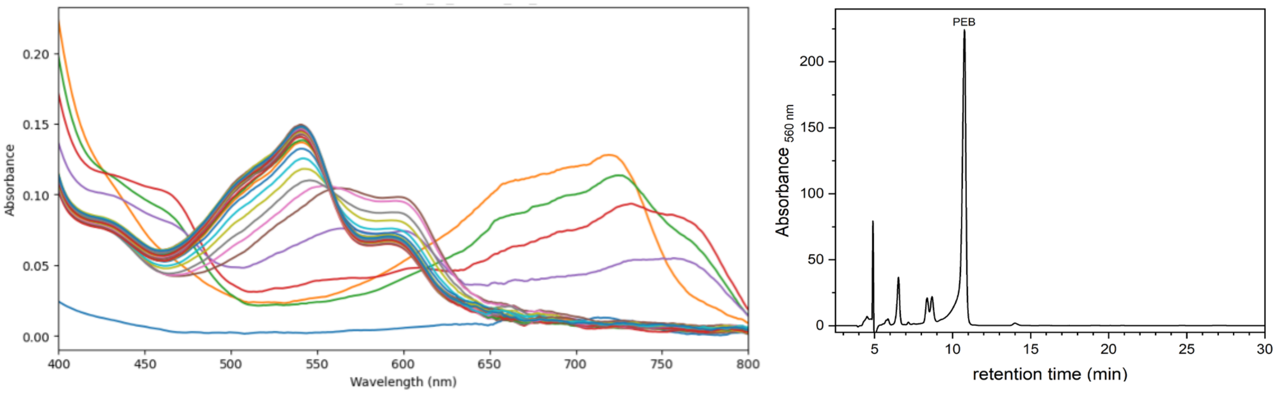


**Figure S7: Activity of wild-type PebS with Te-Fd and PSSM2-Fd.** Time-resolved UV/Vis spectroscopy and HPLC analysis of biliverdin IXα (BV) conversion by wild-type PebS in the presence of either Te-Fd A) or PSSM2-Fd B). Absorbance at 650 nm corresponds to BV, while absorbance at 545 nm indicates the formation of phycoerythrobilin (PEB). Reactions with PSSM2-Fd (B) show higher activity and more efficient PEB formation compared to Te-Fd (A), suggesting functional co-adaptation between PebS and its phage-derived ferredoxin.

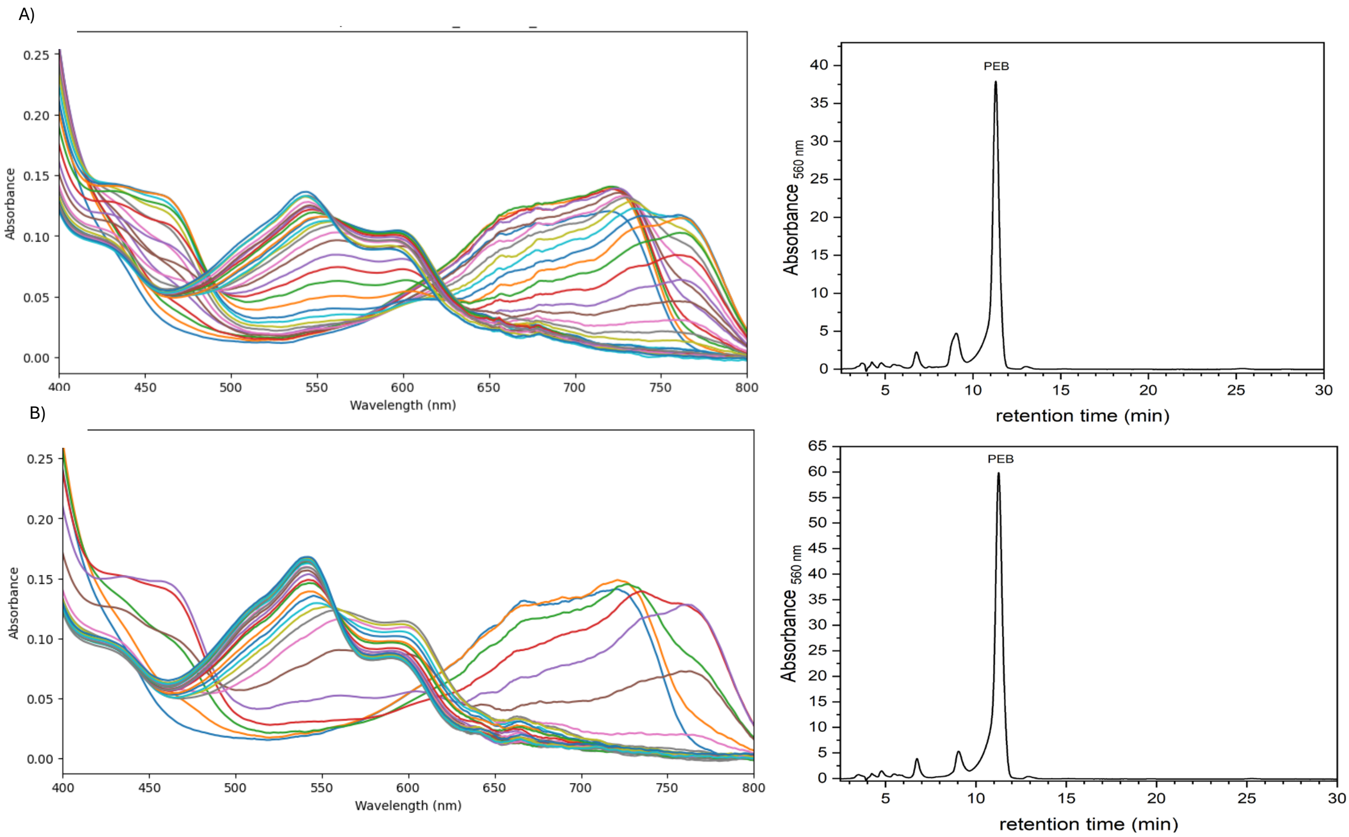


**Figure S8: Time-resolved spectroscopic and chromatographic analysis of biliverdin IXα (BV) conversion to phycoerythrobilin (PEB) by the PebS G210A mutant in the presence of different ferredoxins.** UV/Vis absorbance spectra (left) and corresponding HPLC chromatograms (right) were used to monitor enzymatic activity over 20 minutes, with spectra recorded every 30 seconds. Absorbance at 650 nm indicates the decrease in BV, while absorbance at 545 nm reflects PEB formation. (A) Reactions with T. elongatus ferredoxin (Te-Fd). (B) Reactions with PSSM2-derived ferredoxin (PSSM2-Fd).

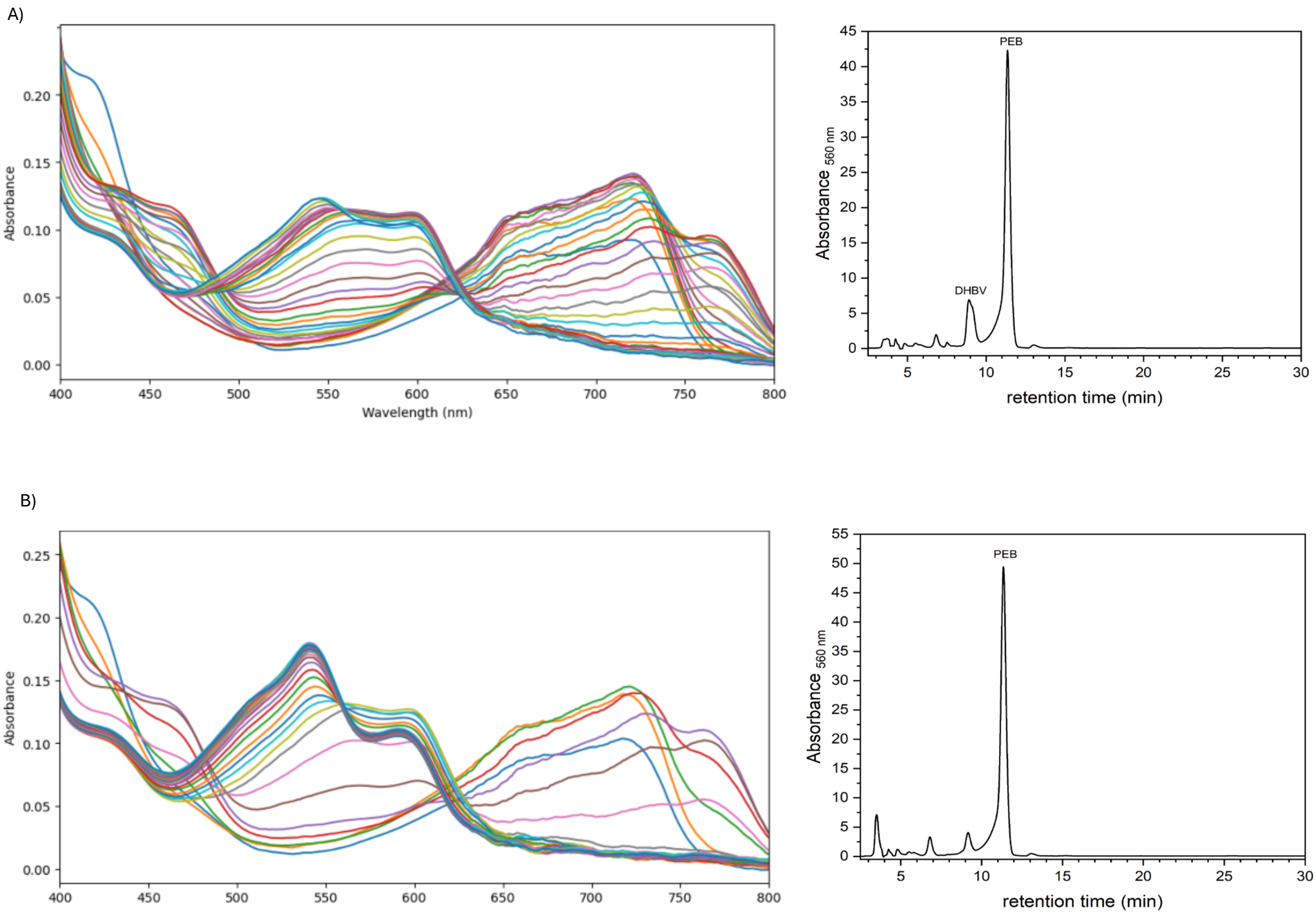


.

**Figure S9: Time-resolved conversion of BV to PEB by the PebS F143Y mutant in the presence of different ferredoxins.** UV/Vis spectroscopy was used to monitor the enzymatic activity of PebS F143Y over 20 minutes, with spectra recorded every 30 seconds. Absorbance at 650 nm indicates the presence of biliverdin IXα (BV), while absorbance at 545 nm corresponds to the formation of phycoerythrobilin (PEB). HPLC chromatograms confirm the identity of the reaction products.A) Reaction with Te-Fd (UV/Vis, left; HPLC, right). B) Reaction with PSSM2-Fd (UV/Vis, left; HPLC, right).

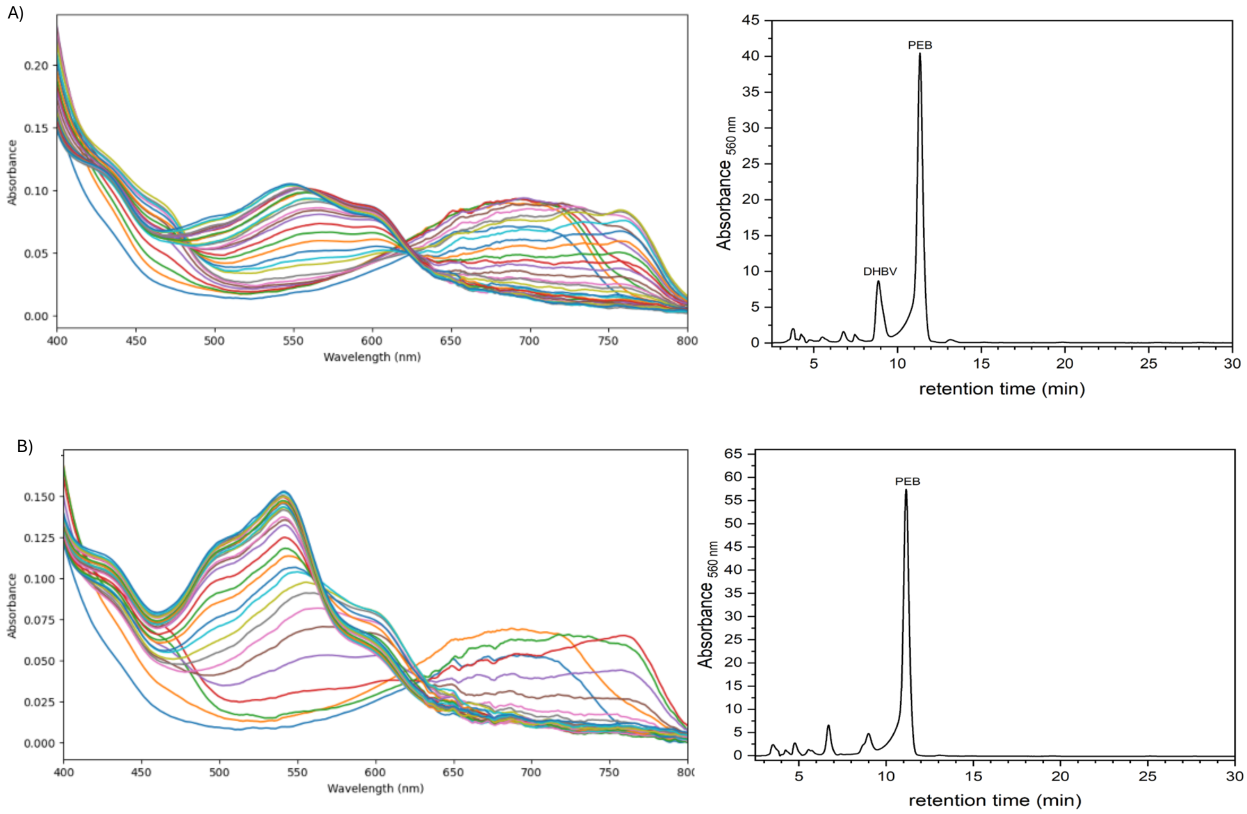


**Figure S10: Time-resolved conversion of BV to PEB by the PebS Y211F mutant in the presence of different ferredoxins.** UV/Vis spectroscopy was used to monitor the activity of PebS Y211F over a 20-minute period, with measurements taken every 30 seconds. Absorbance at 650 nm reflects biliverdin IXα (BV), while absorbance at 545 nm corresponds to phycoerythrobilin (PEB) formation. HPLC analysis confirms the identity of reaction products. A) Reaction with Te-Fd (UV/Vis, left; HPLC, right). B) Reaction with PSSM2-Fd (UV/Vis, left; HPLC, right).

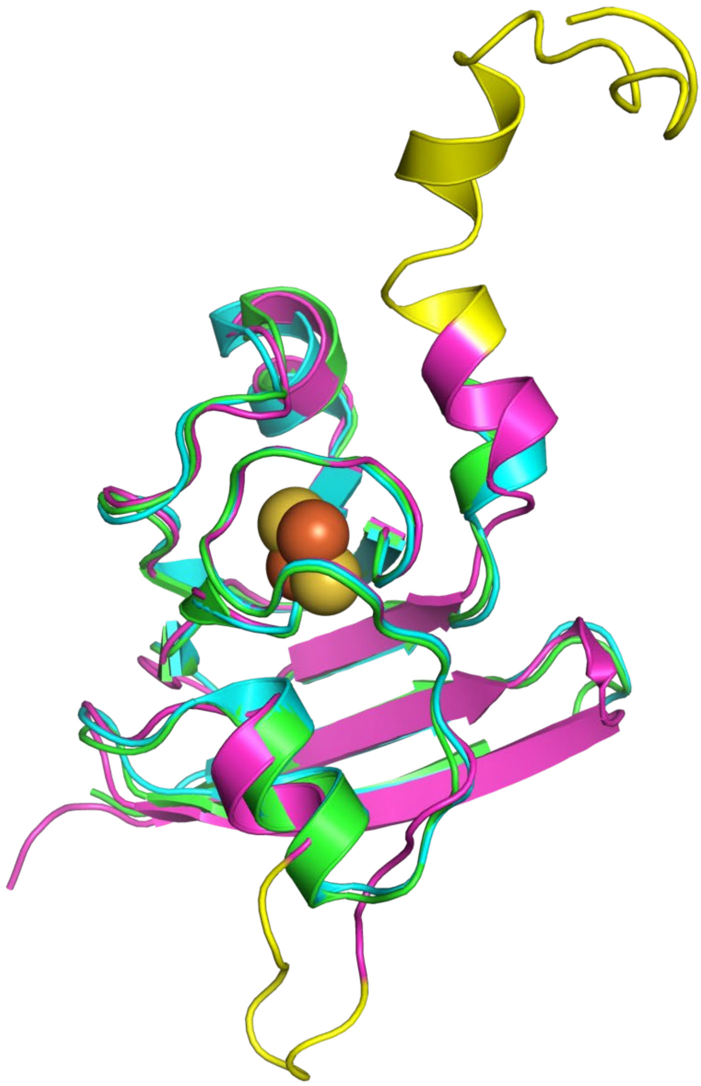

**Figure S11: Structural alignment of ferredoxins.** Shown is a superposition of Te-Fd (PDB ID: 5AUI, green), PSSM2-Fd (PDB ID: 6VJV, cyan) and NATL1A-Fd (Alphafold2 model, purple). Shown is a cartoon representation, atoms of the [2Fe-2S] cluster are shown as spheres. The non-conserved sequence inserts in NATL1A-Fd are colored yellow.
